# 3D association of ER, PR and Her2 at protein levels to aid sub-grouping of breast cancer patients in routine clinical practice

**DOI:** 10.1101/584599

**Authors:** Jiahong Lv, Guohua Yu, Lei Li, Yunyun Zhang, Wenfeng Zhang, Yan Lv, Jiandi Zhang, Fangrong Tang

**Affiliations:** Yantai Quanticision Diagnostics, Inc, a division of Quanticision Diagnostics, Inc at RTP, NC, USA. Yantai, P. R. China.; Dept. of Molecular Pathology, Yuhuangding Hospital, affiliated Qingdao University, Yantai, P. R. China.; Dept. of Pathology, Xintai People’s Hospital, Xintai, P. R. China.; Center for Precision Medicine, Binzhou Medical University, Yantai, P. R. China.

**Keywords:** QDB, 3D, biomarker, absolute, continuous variables, spatial relationship, breast cancer

## Abstract

Treatments of patients with solid tumors frequently rely on the relative and semi-quantitative assessment of tissue protein biomarkers by immunohistochemistry (IHC). The perspective of transition to absolute and quantitative assessment of tissue biomarkers is hindered by lack of a suitable method, especially for Formalin Fixed Paraffin Embedded (FFPE) tissues. In this study, we explored the feasibility of Quantitative Dot Blot (QDB) method as a universal platform to quantitate tissue biomarkers as absolute and continuous variables in FFPE samples by measuring unprecedentedly the protein levels of Estrogen Receptor (ER), Progesterone Receptor (PR), Her2 and Ki67 simultaneously in 1048 FFPE samples. When using measured Her2, ER and PR levels as coordinates to develop 3D scatterplot, we observed a distinct distribution pattern of the samples with natural segregation of three groups as the likely phenotypical projection of known intrinsic subtypes. Thus, we have achieved a major milestone in this transition by identifying the first practice method for daily clinical practice, and one clinical usage in 3D subtyping of samples for prediction and prognosis. This study may serve as basis for a new field of Quantitative Diagnosis where diagnosis, prognosis and prediction are derived from database analysis of protein biomarkers as absolute and continuous variables.

---

As an integral part of clinical diagnosis, immunohistochemistry analysis (IHC) provides valuable information about the localization of biomarkers. Yet, to rely on this method to accurately assess the quantity of tissue biomarkers is technically challenging due to its relative and semi-quantitative nature, let alone other issues including the inherent subjectivity, inconsistency and inter-observer variations. The limitations of IHC method may be used to explain the discordance between microarray-based intrinsic subtyping and IHC-based surrogate assay of breast cancer patients in clinical practice^1–3^.

Hundreds of genes were analyzed in initial microarray analysis of breast cancer tissues to reveal four intrinsic subtypes of breast cancers, including Luminal, Erb2 (Her2 type), basal-like and normal-like (intrinsic subtyping)^4,5^. However, later microarray studies suggested that a few gene modules including ER and Her2 modules may be sufficient for subtyping of breast cancer patients^6–9^. While microarray analysis is limited to retrospective studies only, surrogate assay was developed in clinical practice. This assay was based on the IHC analysis of four protein biomarkers of Estrogen receptor (ER), Progesterone receptor (PR), Ki67 and Her2 to achieve fair but unsatisfactory concordance with intrinsic subtyping^1–3,10^.

Based on these studies, we hypothesized that the intrinsic subtypes of breast cancer patients are ultimately defined by the quantitative protein levels of ER, PR and Her2. The expression levels of multiple genes in the microarray studies^4–8^ may be combined as a quantitative gauge to indirectly reflect the protein levels of ER, PR and Her2, and should be dispensable once the protein levels of these biomarkers are measured quantitatively.

The IHC-based surrogate assay is obviously insufficient in this aspect by its dichotomous classification of protein biomarkers^3–5^. Indeed, studies based on Selected Reaction Monitoring Mass Spectrometry (SRM-MS), the only available method for absolute quantitation of tissue biomarkers in Formalin Fixed Paraffin Embedded (FFPE) samples, showed repeatedly that when Her2 levels were quantitatively measured, there existed as much as over 100 fold variations among those FFPE specimens that tested Her2 positive by IHC and/or FISH analysis^11–15^.

Recently, a high throughput immunoblot method, Quantitative Dot Blot analysis (QDB), was developed in the company^16,17^. This method provides a feasible alternative for absolute quantitation of tissue biomarkers in FFPE samples to SRM-MS method, which so far is limited to only a few proteins ^11–13,15^. We explored QDB method as a universal platform to develop assays using a group of clinically validated antibodies for IHC analysis (IVD or ASR antibody), so that we could test our hypothesis with FFPE samples.

We measured ER, PR, Her2 and Ki67 levels as absolute and continuous variables in 1048 FFPE samples using QDB-based assays (Extended data 1). In this retrospective study, all the samples were collected sequentially and nonselectively as 2X15 μm FFPE slices from local hospitals. These samples were used for preparation of total tissue lysates by deparaffinization and solubilization with lysis buffer. All four biomarkers were measured using the same lysate prepared from 1048 FFPE samples, with intra and inter-CVs below 15% when measured three times, each time in triplicate. To ensure the consistency of the method, the absolute levels of both Her2 and Ki67 of the first 332 samples were measured with two IHC antibodies independently. The correlations of the measured results were all above 0.96 when analyzed with Pearson’s correlation analysis (Extended Data 2a-b).

The distribution of all four biomarkers were shown in Fig 1a-1d. The correlations of QDB results with provided IHC results were shown in Fig 1e-1h. Our results were found to be highly correlated with IHC results for Her2 (ρ=0.58, p<0.0001), ER (ρ=0.60, p<0.0001), PR (ρ=0.63, p<0.0001), and Ki67 (ρ=0.54, p<0.0001) when analyzed with Spearman’s rank correlation analysis. In addition, for ER, PR, and Ki67, when sub-grouping the samples based on their respective IHC scores, the correlations between the sub-group averages of QDB levels and IHC scores increased significantly to r=0.80 for ER, r=0.60 for PR, and r= 0.91 for Ki67 with Pearson’s correlation analysis. We also observed highly similar patterns as reported previously when plotting ER or PR with Her2 as absolute and continuous variables^18^ (Extended Data 3)

**Fig. 1:**
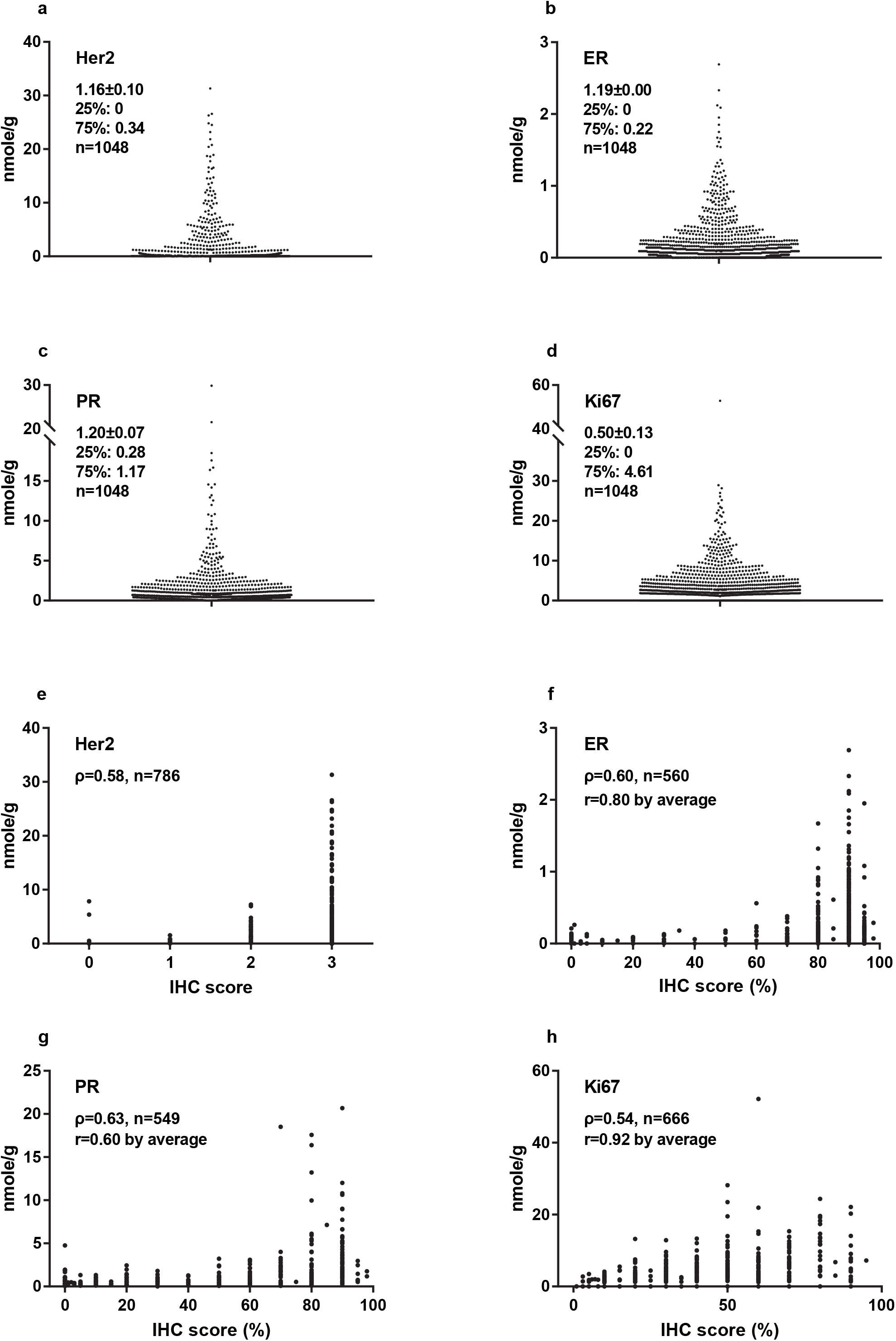
Distribution of Her2, Estrogen receptor (ER), Progesterone receptor (PR) and Ki67 levels as absolute and continuous variables among 1048 breast cancer FFPE samples (1a-1d), and their correlations with IHC scores provided by local hospitals (1e-1h). 1a-1d: The absolute and quantitative levels of Her2, ER, PR, and Ki67 were measured with Quantitative Dot Blot (QDB) method using lysates prepared from 2X15 μm FFPE slices. The average levels of these biomarkers were expressed as mean ± SEM. The 25th and 75th percentiles were also listed for each biomarker respectively. The results were reported as the average of three independent measurements of these four biomarkers, with each measurement in triplicate. 1e-1h: The correlations between QDB and IHC results were analyzed among those samples with available IHC scores using Spearman’s rank correlation analysis with p<0.0001 for all analysis. For ER, PR and Ki67, these samples were sub-grouped based on their IHC scores, and correlation of their subgroup averages with the matching IHC scores were analyzed again with Pearson’s correlation analysis.

The putative relationships among ER, PR and Her2 were investigated by constructing a three-dimensional scatterplot using their protein levels as X, Y and Z coordinates (Fig. 2a). Considering the tissue heterogeneity of breast cancer, it should be emphasized that this relationship is only valid when all the biomarkers are measured from the same lysate.

**Fig. 2:**
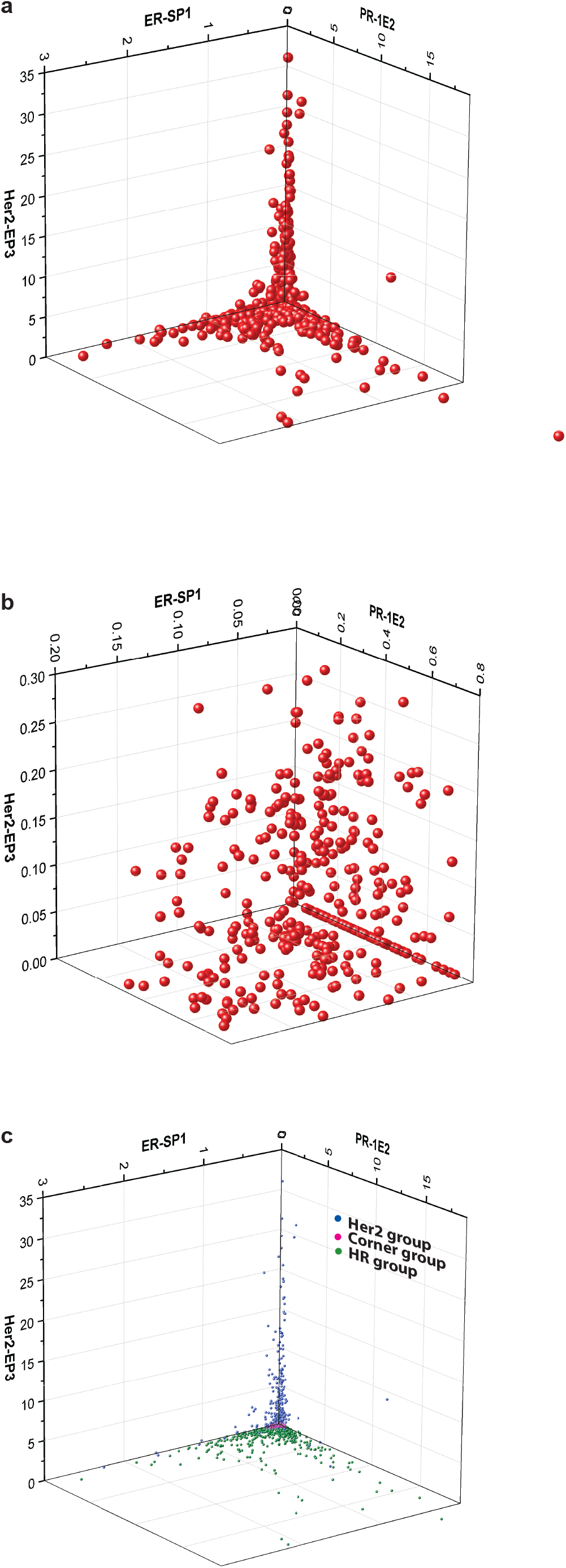
Three-dimensional distribution of 1048 FFPE samples based on the absolute and quantitative levels of ER, PR and Her2. (a) The three-dimensional scatterplot using absolute and quantitative ER, PR and Her2 levels as X, Y and Z axes was created with Origin 9.1 software. Samples were segregated into three groups to resemble a “ball falling from ceiling corner” model as described in the text. (b) The 3D distribution of the samples was narrowed down constantly into a small block with ER<0.2 nmole/g, PR<0.8 nmole/g and Her2<0.3 nmole/g where the samples were found distributed randomly inside. (c) Illustration of HR, Her2 and Corner groups by color in 3D scatterplot.

We observed a pattern we called “balls falling from the ceiling corner”, with three distinctive sub-groups appearing based on the spatial distributions of the samples. We named the first group of samples Hormone receptor (HR) group as they spread flat on the ER-PR plane, representing samples with dominant expression of hormone receptors and minimum Her2 expression. The second group were named Her2 group as they were found wrapping on the Her2 axis, representing samples with strong Her2 expression and minimum hormone receptors expression. The third group accumulated at the intersections of ER, PR and Her2 axes, representing samples lacking strong expressions of ER, PR and Her2. We named this group the corner group. Few samples were found floating in the ER-PR-Her2 space, indicating the lack of samples that strongly expressed all three biomarkers of ER, PR and Her2.

The “balls falling from the ceiling corner” pattern persisted until we narrowed the view into a small block of ER <0.2 nmole/g, PR <0.8 nmole/g and Her2 <0.3 nmole/g, where we began to find samples distributed randomly inside (Fig 2b). Therefore, we used these values as cutoffs to separate samples as inside/outside the range with 756(72.1%)/292(27.9%) for ER; 688(65.7%)/ 359(34.3%) for PR and 777(74.2%)/271(25.8%) for Her2. Consequently, we were able to assign 412 samples into HR group (39.4% of total 1048 samples, including 168 samples with both ER and PR outside of the range), 271 into Her2 group (25.8%), and 365 into corner group (34.8%).

Interestingly, we identified 211 out of 271 (77.9%) samples from Her2 group within both the ER and PR ranges. For the remaining 60 samples, 49 out of 60 (83.3%) were within either the ER or PR range. In other words, we were able to identify 260 out of 271 (96.3%) in the Her2 group by limiting either ER <0.2 nmole/g or PR <0.8 nmole/g. Among the 11 outliers, 8 samples were at the vicinity of ER or PR cutoffs. The only three exceptions were with their Her2, ER and PR levels at (3.57, 0.45, 1.13), (1.74, 0.69, 0.97) and (0.42, 0.40, 1.19). These samples were also the only ones with medium to strong expressions of all three biomarkers, in agreement with our observations of few samples floating in the ER-PR-Her2 space.

The Ki67 levels of these three groups were evaluated in Fig.3a. We found the averaged Ki67 levels were 3.80±6.17 nmole/g, 4.28±3.80 nmole/g, and 2.84±3.10 nmole/g for Corner, Her2 and HR groups respectively. There were statistical differences between corner group and HR group (p=0.0058), between Her2 group and HR group (p<0.0001), but not between corner group and Her2 group when Ki67 levels were analyzed with unpaired student t-test. The corner group was also the most heterogeneous group concerning Ki67 levels. Its median level was the lowest among the three groups (1.71 vs 3.03 for Her2 group and 2.08 for HR group), yet the top 1% of samples by Ki67 expression were all within this group.

**Fig. 3:**
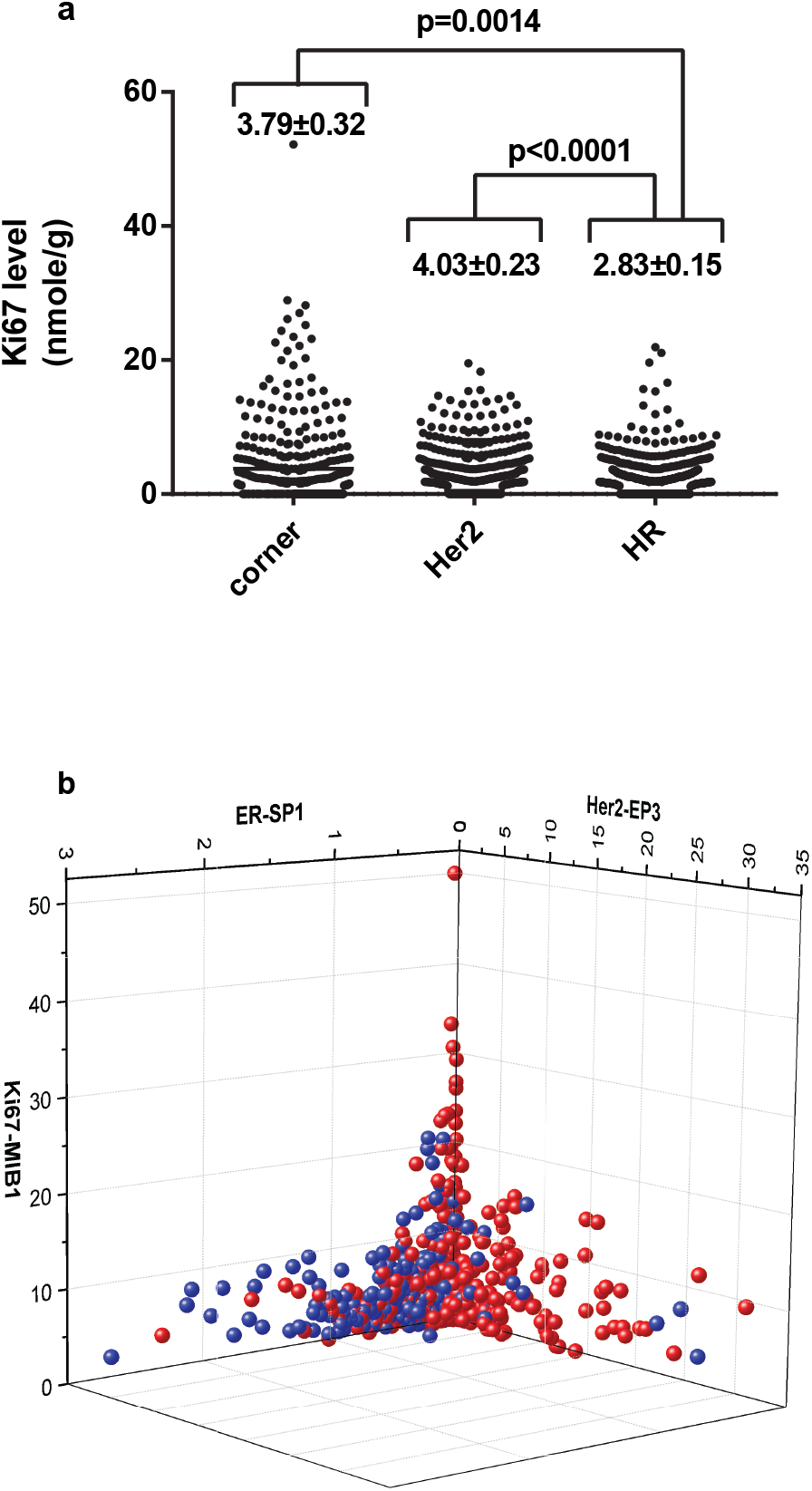
Evaluation of Ki67 levels of samples grouped by their spatial distribution. (a) Comparison of Ki67 levels among three spatial groups of HR, Her2 and Corner groups. The absolute and quantitative Ki67 levels of all 1048 samples were measured with QDB method using the same lysates for ER, PR, and Her2 measurements. The samples were grouped into HR, Her2 and Corner groups based on the observed cutoffs at Fig. 2b (Her2 group, Her2≥0.3 nmole/g; HR group, ER≥0.2 nmole/g and/or PR≥0.8 nmole/g; corner group, ER<0.2 nmole/g, & PR<0.8 nmole/g & Her2<0.3 nmole/g). The results were analyzed using unpaired two-tailed student t-test. (b) The spatial distribution of samples using ER, Her2 and Ki67 levels as X, Y and Z axes. Those samples with PR level <0.8 nmole/g were arbitrarily set to the color red, while those with PR level ≥0.8 nmole/g were set to the color blue. We observed that those samples with the highest Ki67 levels were exclusively red around the intersection of ER and Her2, suggesting a lack of ER, PR, and Her2 expression.

When observing the 3D scatterplots of ER-Her2-Ki67 (Fig 3b), ER-PR-Ki67 and PR-Her2-Ki67 together (Extended Data 4a & b), we found that samples with highest Ki67 levels were at the intersection of ER, PR and Her2. We managed to include PR information in the 3D scatterplot of ER-Her2-Ki67 by assigning samples with PR <0.8 nmole/g as red, and PR≥0.8 nmole/g as blue in Fig. 3b. Samples with highest Ki67 levels were found exclusively in red in this picture. In addition, samples within the top 10th percentile by Ki67 levels were either within the ER and/or PR range (91%), or at the vicinity of these cutoffs (9%).

The clinical relevance of the suggested block of ER <0.2 nmole/g, PR <0.8 nmole/g and Her2 <0.3 nmole/g was explored next. We hypothesized that these cutoffs might correspond to the cutoff values to differentiate negative samples from positive ones in clinical practice. For this purpose, the recommendations from ASCO/CAP were followed to differentiate samples into Her2+ from Her2-based on IHC results, and Receiving Operative Characteristic (ROC) analysis was performed with measured Her2 levels. As expected, we confirmed Her2 at 0.3 nmole/g as the optimized cutoff value to achieve the best sensitivity and specificity at 84.8% and 97.6% respectively with IHC results, with overall concordance rate at 93.6% with IHC analysis (Fig. 4a & 4b).

**Fig. 4:**
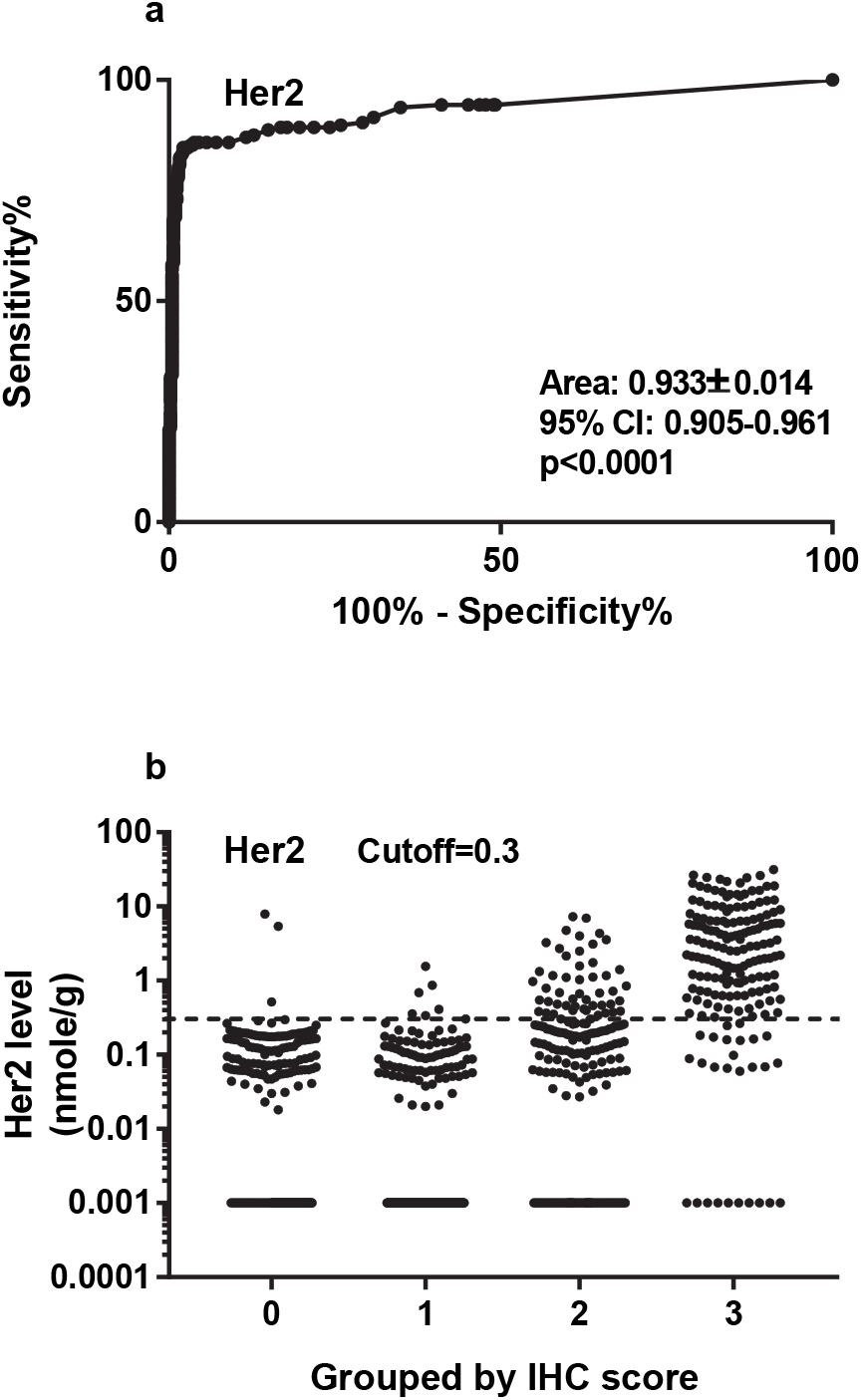
Evaluating the sensitivity and specificity of QDB method using Receiver Operative Characteristics (ROC) analysis with provided Her2 IHC scores from local hospitals. (a) Samples with provided IHC scores were grouped as negative (IHC scores of 0 and 1+) or positive (IHC scores of 3+), and were used for Receiver Operative Characteristics (ROC) analysis using GraphPad Software. We were able to achieve Area Under the Curve (AUC) at 0.933±0.014, 95%CI at 0.905~0.961 (p<0.0001). Using 0.3 nmole/g as the cutoff, we achieved sensitivity at 84.8% and specificity at 97.6%. The concordant rate was at 93.6% (560 out of 598, samples with a Her2 score of 2+ were excluded from analysis) with IHC results. (b) To better illustrate the effectiveness of this cutoff (indicated by the dashed line) at separating negative samples from positive ones, samples were plotted in log scale and grouped based on their respective IHC scores. All those samples with their Her2 levels measured as 0 were arbitrarily set as 0.001 nmole/g to avoid being omitted in the log scale plot.

However, when using the recommended IHC score at 1% by ASCO/CAP as cutoff to differentiate ER and PR into negative and positive groups, we achieved better sensitivity and specificity using 0.045 nmole/g and 0.47 nmole/g as cutoffs for ER and PR than the observed 0.2 nmole/g and 0.8 nmole/g for ER and PR (data not shown). Yet, we failed to observe any distributional differences graphically around these optimized cutoff values in the 3D scatterplot. In addition, the proposed 1% as cutoff value for IHC analysis of ER positivity is not without dispute^19^. In ELISA analysis, the FDA-approved cutoff value for ER is at 0.15 nmole/g^20^.

The natural segregation of breast cancer samples in 3D scatterplot supports our hypothesis that the intrinsic subtyping of breast cancer patients may be determined by the quantitated protein levels of ER, PR and Her2. Admittedly, we are still unclear about how each intrinsic type is projected in this 3D plot. Future comparative studies with frozen tissues are needed to answer these questions. Nonetheless, our results signaled the need for the transition to absolute quantitation of tissue biomarkers in clinical diagnosis.

In addition, we may also identify a novel predictor model with the 3D relationship among biomarkers at protein level (3D subtyping). Considering biological functions are mainly carried out at protein level, 3D subtyping should offer a unique clinical benefit over those at genetic level. In addition, we expect the application of 3D subtyping with other cancer types in the future.

In summary, our results emphasized the need for a transition to absolute quantitation of tissue biomarkers by demonstrating the natural spatial sub-grouping of breast cancer patients by ER, PR and Her2 protein levels. We also demonstrated QDB to be an effective method to meet this need in daily clinical practice by adopting this method to measure ER, PR, Her2 and Ki67 in 1048 FFPE samples for the first time.

Our study may affect the field of clinical diagnosis in multiple aspects. First, the feasibility of QDB method with FFPE specimen allows us to access the rich reservoir of FFPE specimens worldwide to unprecedentedly generate knowledge about the clinical relevance of tissue biomarkers and their putative associations through population studies; second, it will also speed up the process of identification and verification of new biomarkers in clinical diagnosis; and third, the existence of hundreds of IVD or ASR antibodies would allow us to extend our studies to a wide spectrum of solid tumors expediently. These studies may lead to a new field of quantitative diagnosis, where the clinical decisions are made from population studies of multiple biomarkers as absolute and continuous variables.

## MATERIALS AND METHODS

### Human subjects and human cell lines

A total of 1048 Formalin Fixed Paraffin Embedded (FFPE) breast cancer tissue specimens were provided sequentially and non-selectively together with IHC scores of four biomarkers from some of the specimens from Yantai Yuhuangding Hospital, and Xintai People’s Hospital from P. R. China. All the samples were obtained in accordance with the Declaration of Helsinki and approved by the Medical Ethics Committees of each institutions respectively.

MCF-7 and BT474 cell lysates were obtained from the Cell Bank of Chinese Academy of Sciences (Shanghai, China), and used as controls for all four biomarkers.

### General reagents

Recombinant human Her-2/ErbB2 protein was purchased from Biological Inc. ER, PR and Ki67 recombinant proteins were purified in the house. QDB plate was purchased from Quanticision Diagnostics, Inc (RTP, NC, USA). Ventana anti-Her2/neu(4B5) rabbit monoclonal primary antibody and anti-PR (1E2) rabbit monoclonal primary antibody were purchased from Roche Diagnostics GmbH. Rabbit anti-ER (SP1) antibody was purchased from Abcam Inc. Rabbit anti-Her2 antibody (EP3) and mouse anti-Ki67 (MIB1 & UMAB107) were purchased from ZSGB-BIO (www.zsbio.com, Beijing, China). HRP labeled Donkey Anti-Rabbit IgG secondary antibody was purchased from Jackson Immunoresearch lab (West Grove, PA, USA). BCA total protein quantification kit was purchased from Thermo Fisher Scientific Inc (Calsband, CA, USA).

### Purification of protein standards for ER, PR and Ki67

DNA sequences corresponding to the 1162-1254AA of human MKI67 (NCBI #: NM_002417.4), 455-595AA of human ER-α (NCBI #: NM_000125.3), and 310-417 AA of human PR isoform B (NCBI#: NP000917.3) were inserted into pET-32a (+) expression vector respectively and expressed in E.coli BL21(DE3) competent cells.

The cells were induced with 1mM IPTG, and total bacterial lysates were extracted in 10ml binding buffer(20mM sodium phosphate, 500mM NaCI, 20mM imidazole, PH 7.4) before they were loaded onto a high affinity Ni^2+^ column pre-equilibrated with 10ml binding buffer. The recombinant protein was eluted with 3ml elution buffer (20mM sodium phosphate, 500mM NaCI, 250mM imidazole, PH 7.4), and dialyzed in PBS (PH 7.4) at 4°C overnight. The purity of the protein was examined by a 12% SDS-PAGE gel, and the purified protein was stored at −80°C in small aliquot with 20% glycerol.

### Preparation of FFPE tissue and cell lysates

Two whole tissue slices (2X15μm) from FFPE blocks were put into 1.5ml Eppendorf tubes, and deparaffinized before they were solubilized using solubilization buffer(50mM Tris-HCl, 10 mM EDTA, 1% Tween 20, 10% glycerol, pH 9.9). Cells (MCF-7 & BT474 cells) were lysed in the lysis buffer (50mM HEPES, 137mM NaCl, 5mM EDTA, 1mM MgCl_2_, 10mM Na_2_P_2_O_7_, 1%TritonX-100, 10% glycerol, pH7.6) with protease inhibitors (2μg/ml Leupeptin, 2μg/ml Aprotinin, 1μg/ml pepstatin, 2mM PMSF, 2mM NaF). The supernatants were collected after centrifugation and the total amount of protein was measured using BCA protein assay kit by following manufacturer’s instructions.

### QDB analysis

Sample pools were prepared by mixing tissue lysates from four FFPE tissue specimens with an IHC score of 3+ for Her2, and IHC score of 90% for ER, PR and Ki67 to define the linear range of QDB assay respectively. The pooled lysates were serially diluted side by side with the recombinant proteins for defining the standard curves of QDB analysis.

The QDB process was described elsewhere with minor modifications^16,17^. In brief, the final concentration of the FFPE tissue lysates was adjusted to 0.25 μg/μl for Her2 and Ki67 and 0.125 μg/μl for ER and PR, and 2 μl/unit was used for QDB analysis in triplicate. The QDB plate was then dried for one hour at RT, soaked in transfer buffer for 10s, rinsed once with TBST, and then blocked in 4% non-fat milk for an hour. Next, it was put into a 96-well microplate with 100μl primary antibody (for clone EP3, 1:1500 in blocking buffer; for clone 4B5, 1:10 in PBS; for clone SP1, 1:250 in blocking buffer; for clone 1E2, 1:8 in PBS; for clone MIB-1, 1:1000 in blocking buffer)), and incubated overnight at 4°C. Afterward, the plate was rinsed twice with TBST and washed 3X10mins. It was then incubated with either a donkey anti-rabbit or donkey anti-mouse secondary antibody for 4 hours at RT, rinsed twice with TBST, and washed 4X10mins. Finally, the QDB plate was inserted into a white 96-well plate pre-filled with 100μl/well ECL working solution for 3mins. The chemiluminescence signal of the combined plate was quantified by using the Tecan Infiniti 200pro Microplate reader with the option “plate with cover”.

For the 1048 FFPE samples, each sample was measured three times, each time in triplicate. The consistency of the experiments was also ensured by including BT-474 and MCF-7 cell lysates with pre-documented biomarker levels in all the experiments respectively. The result was considered valid when the calculated biomarker levels of control cells were within 10% of the pre-documented biomarker levels. The absolute biomarker levels were determined based on the dose curve of protein standard, with those samples with chemiluminescence reading of less than 2 fold over blank were defined as non-detectable, and entered as 0 for data assay.

### Data analysis

All the data were presented as Mean±SEM. All the 3D analyses of biomarkers were performed using Origin pro 9.1 software from Originlab Corp (Northampton, Massachusetts, USA). All the statistical analyses, including the unpaired two-tailed student t test, were performed with GraphPad Prism software version 7.0 (GradphPad Software, USA). P value <0.05 was considered statistically significant.

## Code Availability statement

Not applicable.

---

**Extended Data 1.**
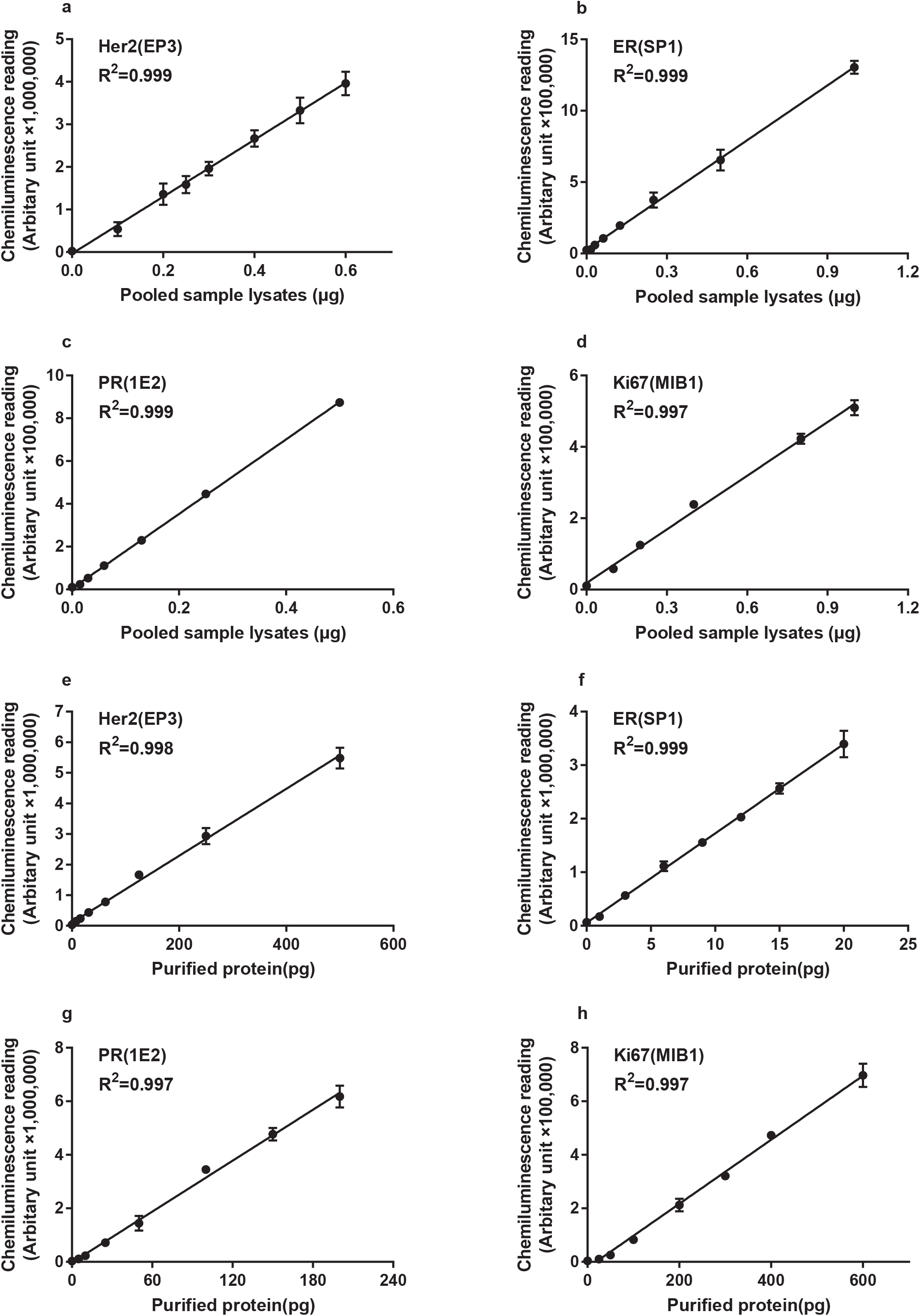
Defining the linear ranges of QDB method for measuring Her2, Estrogen receptor (ER), Progesterone receptor (PR) and Ki67 levels using validated IHC antibodies as indicated in the figure. Fig. 1a-1d: Pooled samples prepared from 2X15 μm FFPE slices obtained from 4 patients testing positive based on IHC analyses of these biomarkers were used to define the linear range of QDB method for these biomarkers respectively. Pooled samples were serially diluted and supplemented with IgG-free BSA at a final amount of 0.5 μg/unit for Her2 and Ki67, and 0.25 μg/unit for ER and PR. The chemiluminescence signals were captured with a Tecan microplate reader, and used for linear regression analysis with the matching amount of total protein lysates used in the QDB analysis. The linear range of the analysis was defined as where the coefficient of determination (R^2^) was above 0.99. Fig. 1e-1h: Purified recombinant protein, either obtained commercially (Her2), or purified in the house (ER, PR and Ki67) were loaded at the amount indicated in the figure legends, and supplemented with IgG-free BSA to match the final loading amount for sample analysis at 0.25 μg/unit for ER and PR, and 0.5 μg/unit for Her2 and Ki67. The chemiluminescence signals were captured with a Tecan microplate reader, and used for linear regression analysis with the matching amount of protein standard used in the QDB analysis. The linear range of the analysis was defined as the region where the coefficient of determination (R^2^) was above 0.99.

**Extended Data 2:**
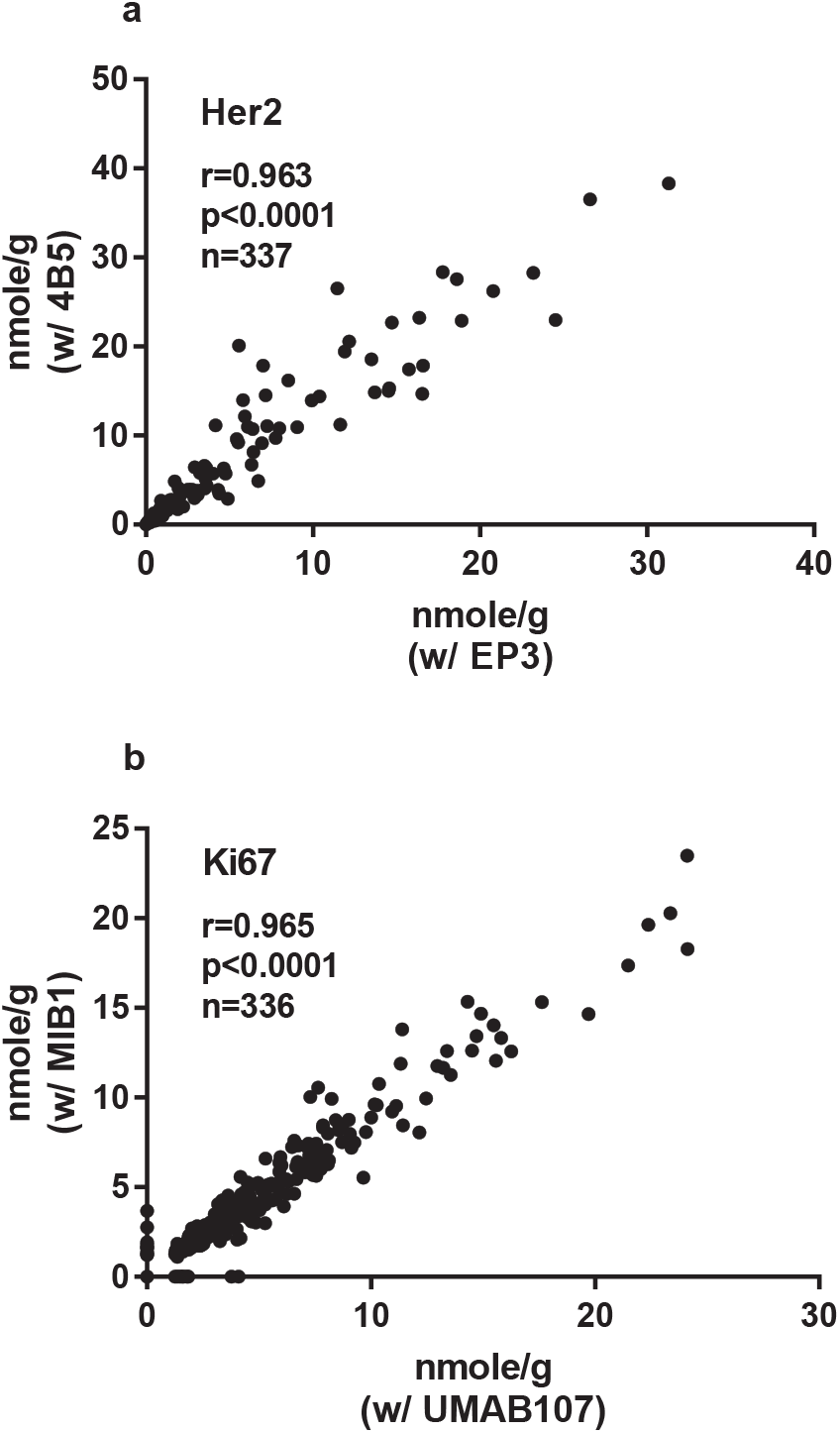
Consistency of the QDB measurement of Her2 (a) and Ki67 (b) levels using two independent validated IHC antibodies respectively. (a) Total lysates prepared from 2X15 μm FFPE slices were used for QDB measurements of Her2 levels at 0.5 μ g/sample with EP3 and 4B5 antibodies respectively, as described in Materials and Methods. The correlation between Her2 levels from EP3 and 4B5 measurements were analyzed with Pearson’s correlation analysis in 332 samples. P<0.0001. (b) The same lysates from (a) were also used for measurements of Ki67 levels using MIB1 and UMAB107 respectively in 332 samples. The correlation between Ki67 levels from MIB1 and UMAB107 measurements were analyzed using Pearson’s correlation analysis. P<0.0001.

**Extended Data 3.**
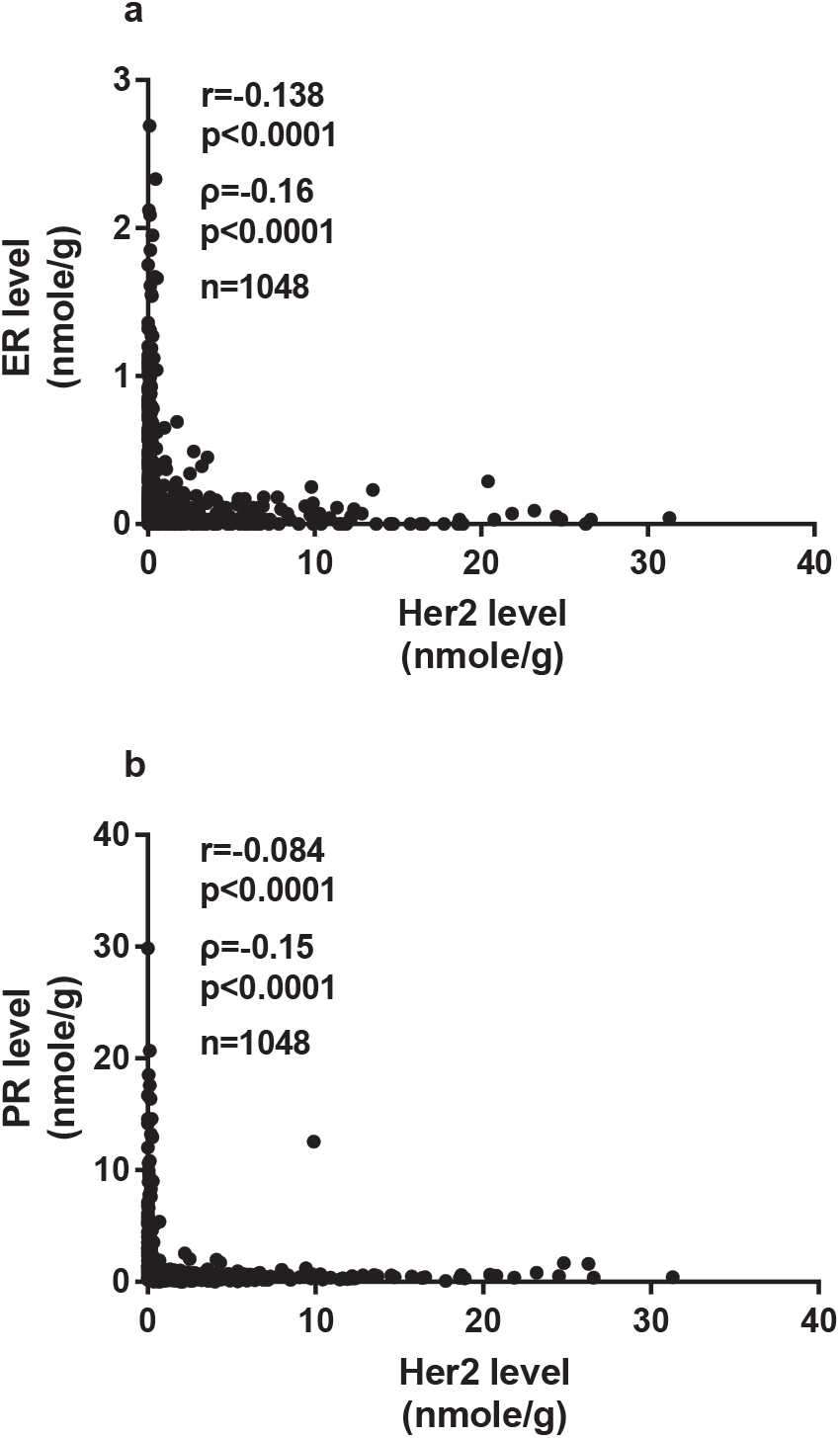
The relationship between Her2 and ER (a) or PR (b) by absolute and quantitative protein levels. The correlation between these factors was evaluated with both Pearson’s correlation analysis and Spearman’s rank correlation analysis using GraphPad Software.

**Extended Data 4:**
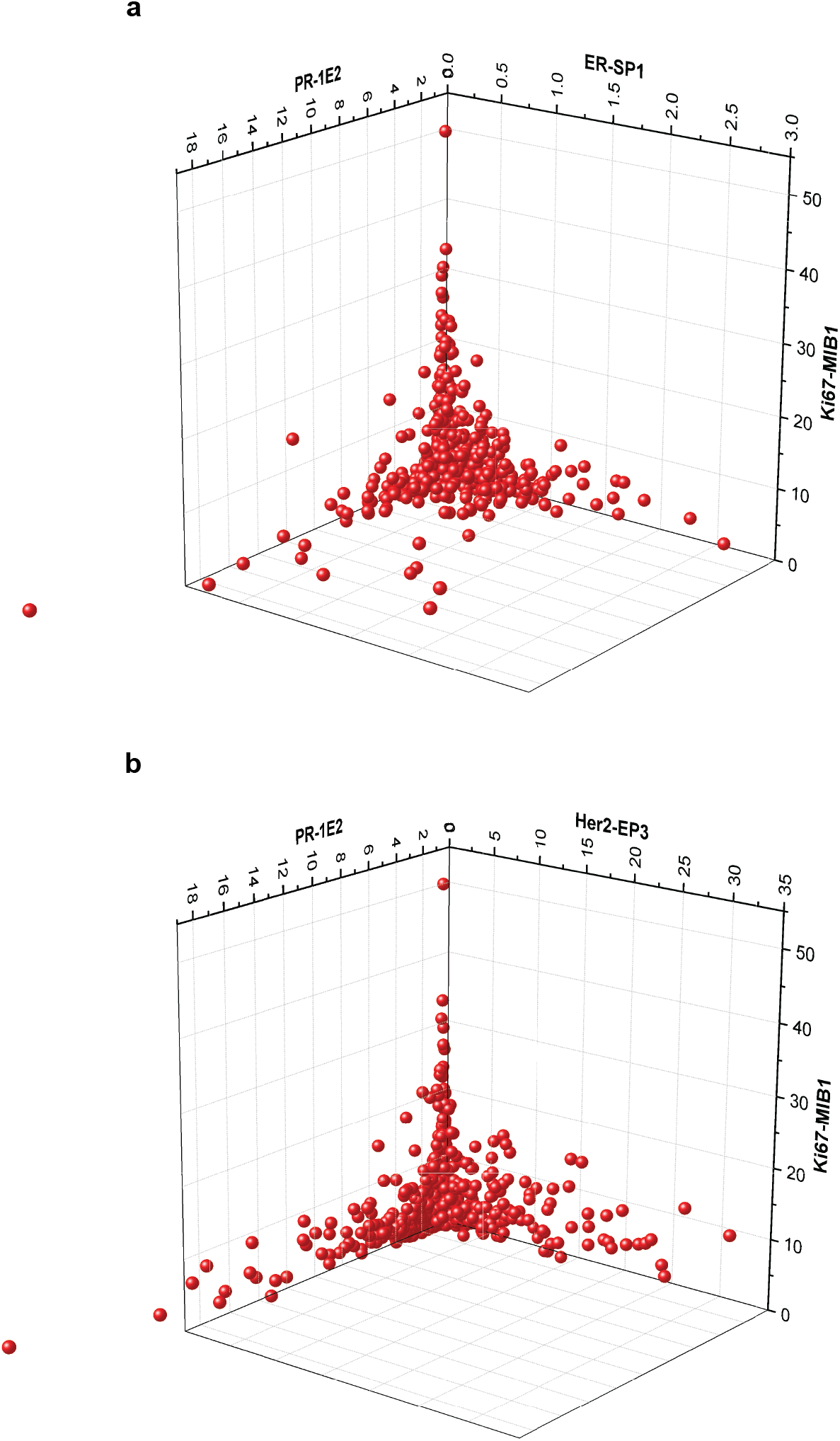
The three-dimensional relationships among PR, ER and Ki67 (a) and PR, Her2 and Ki67 (b) at protein level. The results were plotted in a 3D scatterplot using Origin Pro 9.1.

